# The impact of prenatal alcohol and synthetic cannabinoid exposure on behavioral adaptations in adolescent offspring and alcohol self-administration in adulthood

**DOI:** 10.1101/2023.10.09.561571

**Authors:** Laura C. Ornelas, Eric W. Fish, Jacob C. Dooley, Megan Carroll, Scott E. Parnell, Joyce Besheer

## Abstract

Prenatal exposure to alcohol or cannabinoids can produce enduring neurobiological, cognitive, and behavioral changes in the offspring. Furthermore, prenatal co-exposure to alcohol and cannabinoids induces malformations in brain regions associated with reward and stress-related circuitry. This study examined the effects of co-exposure to alcohol and the synthetic cannabinoid (SCB) CP55,940 throughout gastrulation and neurulation in rats on basal corticosterone levels and a battery of behavioral tests during adolescence and alcohol self-administration in adulthood. Importantly, we find that prenatal alcohol exposure (PAE) caused lower baseline corticosterone levels in adolescent males and females. Co-exposure to alcohol + CP produced hyperactivity during open field test in males, but not females. During the two-bottle choice alcohol-drinking procedure, prenatal cannabinoid exposed male and female adolescent rats drank more alcohol than their vehicle-exposed controls. In adulthood, female rats treated with prenatal cannabinoid exposure (PCE), showed an overall total increase in alcohol intake during alcohol self-administration; but this was not found in males. When the reinforcer was changed to a 1% sucrose solution, male rats exposed to PCE, showed a reduced self-administration compared to vehicle-exposed males, potentially indicative of an anhedonic response. This lower self-administration persisted when 20% alcohol was reintroduced to the sucrose solution. Lastly, following an abstinence period, there were no changes due to prenatal drug exposure in either males or females. Overall, these data suggest lasting consequences of prenatal alcohol and cannabinoid exposure during adolescence and adulthood in male and female rats.

## 1. Introduction

The prevalence of marijuana use in the United States amongst pregnant women has been increasing, specifically over the last 10 years. According to a National Survey on Drug Use and Health in 2018, approximately 9.8% of women reported marijuana use before pregnancy, 4.2% during pregnancy and 5.5% after pregnancy (SAMHSA, 2019). Clinical research has shown that children exposed to prenatal cannabis exhibit altered emotional, behavioral, and cognitive development (Huizink, 2014). Considering the significant risk factors associated with prenatal marijuana exposure, usage of marijuana products during pregnancy is a public health concern that needs to be addressed though continued education, prevention, and research on the effects of marijuana during pregnancy.

Legalization of marijuana for both medical and recreational purposes, as well as the marketing of cannabis-derived cannabinoid products, including synthetic cannabinoids, may lead to the perception that these are low-risk substances safe to use during pregnancy (Chang et al., 2019). Commonly known as ‘Spice’ or ‘K2’, synthetic cannabinoids are compounds that work as full agonists at the cannabinoid CB1 receptor, with binding affinities of 4-5 times higher (Elsohly, Gul, Wanas, & Radwan, 2014) and potencies of 40–660 times higher than THC (van Amsterdam, Brunt, & van den Brink, 2015). There have been a number of adverse side effects associated with the use of synthetic cannabinoids, including cardiovascular effects (i.e. tachycardia, hypertension, arrhythmia) (Tait, Caldicott, Mountain, Hill, & Lenton, 2016), nausea, vomiting, seizures, cognitive deficits, and memory loss (Radhakrishnan, Wilkinson, & D’Souza, 2014; Tait et al., 2016). In addition to the physical health effects, preclinical studies have shown severe brain developmental consequences associated with prenatal synthetic cannabinoid exposure in chickens (Psychoyos, Hungund, Cooper, & Finnell, 2008), mice (Gilbert et al., 2016) and rats (Mereu et al., 2003). While examining later life behavior, prenatal cannabis use has also been found to have long-term effects on diminished verbal and memory skills (Day et al., 1994), increased impulsivity and hyperactivity (Richardson, Ryan, Willford, Day, & Goldschmidt, 2002) and decreased attention and concentration during childhood and adolescence (Fried & Smith, 2001; Martínez-Peña et al., 2021). Consequences of use also affect adult functioning including substance use, memory deficits and cognition, motivation and behavioral regulation (De Genna, Willford, & Richardson, 2022; Zamberletti & Rubino, 2022), as well as altered neural functioning during visuospatial working memory (Smith, Fried, Hogan, & Cameron, 2006). Therefore, while there are limited clinical studies showing the detrimental effects of prenatal synthetic cannabinoids, the risk factors associated with prenatal marijuana exposure are well known; and considering synthetic cannabinoids have stronger binding affinity and potency compared to THC, synthetic cannabinoid use during pregnancy may lead to severe impairments *in utero*.

Co-use of cannabis products with other drugs, including alcohol, is common and is associated with greater risk for negative health outcomes (Coleman-Cowger, Schauer, & Peters, 2017; Metrik & Patel, 2022; Yurasek, Aston, & Metrik, 2017). Between 2015-2018, a national survey on drug use and health showed about 10% of pregnant women had at least one alcohol drink in the past 30 days and of those using alcohol, 40% also used other substances most often tobacco and marijuana (England et al., 2020). Considering the dangers of prenatal cannabis use alone, the combination with alcohol can lead to significantly greater neurodevelopmental effects, as observed in animal models (for review see Rouzer, Gutierrez, Larin, & Rajesh, 2023). Fish et al. 2019, showed prenatal exposure to cannabinoids (THC, CP 55,940 or HU-210) combined with alcohol produced significant eye and face malformations in fetal mice, as well as widespread brain structural changes (Fish et al., 2019). In addition, examination of zebrafish embryos following exposure to cannabinoids and alcohol showed significant defects to midbrain/hindbrain boundary and eye formation (Fish et al., 2019). Furthermore, the physical impairments of co-use of alcohol and cannabinoids during pregnancy can lead to long-term behavioral and neurological disturbances, such as increased risk of neurodevelopmental, cognitive, and/or neuropsychiatric disorders in the offspring (Cook, 2020). However, there is a general lack of knowledge regarding consequences of prenatal co-exposure to alcohol and synthetic cannabinoids into adolescence and adulthood.

The current study took a systematic approach to examine the consequences of prenatal alcohol and synthetic cannabinoid co-exposure on physiological and behavioral adaptations in adolescence and adulthood in male and female rats. Understanding the consequences of prenatal drug exposure on adolescent behavior is vital, considering this is an important developmental period in which individuals engage in risk taking behaviors and are more vulnerable to developing addictive behaviors. Specifically, we investigated the effects of co-exposure during the gastrulation and neurulation developmental stages (gestational day [GD] 8-11) to alcohol and the synthetic cannabinoid (SCB) CP 55,940 on exploratory and anxiety-related behaviors and home cage alcohol drinking during adolescence and alcohol self-administration during adulthood.

## 2. Materials and Methods

### 2.1. Animals

Female Long-Evans rats approximately 10-12 weeks old were mated to males for one hour at the beginning of the light phase of a 12:12 h light:dark schedule. When a successful copulation occurred, designated as GD 0, mated females were single-housed in ventilated cages (Tecniplast, West Chester, PA) with *ad libitum* food and water. Rats were maintained in a temperature and humidity controlled colony with a 12-hour light/dark cycle (lights on at 07:00). All experiments were conducted during the light cycle. Animals were under continuous care and monitoring by veterinary staff from the Division of Comparative Medicine at UNC-Chapel Hill. All procedures were conducted in accordance with the NIH Guide to Care and Use of Laboratory Animals and institutional guidelines.

### 2.2. Synthetic Cannabinoid and Alcohol Teratogens

The prenatal treatment protocol and experimental timeline are illustrated in Figure 1. In order to characterize the effects of prenatal alcohol or SCB or combination of alcohol and SCB exposure in rats, on GD 8-11 (a critical window encompassing the gastrulation and neurulation developmental stages), pregnant Long Evans rats received drug treatments. Ethanol (20% v:v) was given orally via gavage through a 18 gauge stainless steel feeding tube at a dose of 2 g/kg. CP 55,940 (-)-*cis*-3-[2-hydroxy-4-(1,1-dimethylheptyl)phenyl]-*trans*-4-(3-hydroxypropyl)cyclohexanol, Sigma Aldrich, St. Louis, MO) was suspended in a solution of 5% ethanol, 5% alkamuls EL 620 (Rhodia, Cranberry, NJ), and 90% lactated Ringer’s solution immediately before intraperitoneal injection through a 25 gauge needle at a volume of 1ml/kg body weight. Pregnant dams were treated with two daily injections separated by 4 h. The groups were as follows: 1) Water + Vehicle, 2) twice daily water + once daily 0.3 mg/kg CP 55,940, 3) twice daily EtOH (20%) IG + once daily vehicle and 4) twice daily EtOH (20%) IG + once daily 0.3 mg/kg CP 55,940. Twice daily water and EtOH IG injections were 4 hours apart. The dams were then left undisturbed until birth. Once the pups were born, litters were culled to 10/litter on postnatal day (PND) 2. On PND 21, the pups were weaned and double housed with same sex littermates. 1-2 rats were used from the same litter for each sex per treatment groups. Table representing the sample size for each group can be found in Figure 1.

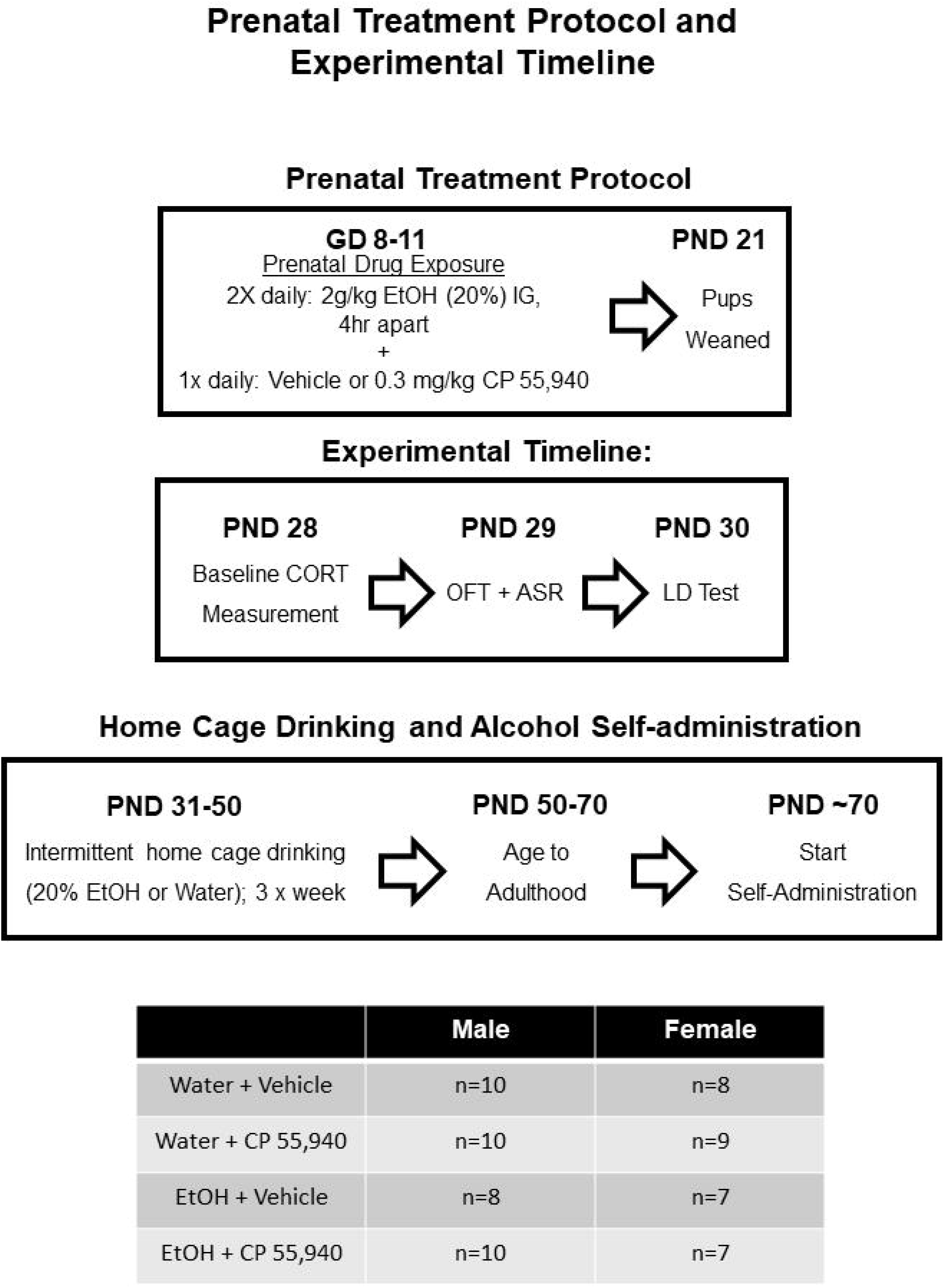

#### Basal Corticosterone Levels

On PND 28 (early adolescence), tail blood was collected (2 hr into the light cycle) into heparinzed tubes and centrifuged at 4°C for 5 minutes at 2000 rcf for analysis of plasma corticosterone levels. Following the completion of blood collection, each rat was returned to the homecage. Approximately 20-30 µL of plasma was collected and stored at −80°C until analysis in duplicate using DetectX^®^ Corticosterone Enzyme Immunoassay Kit (ArborAssays, Ann Arbor, MI) per the manufacturer’s instructions.

#### Open Field Test

On PND 29, rats were placed in an open field for monitoring of general locomotor activity for 20 min. Open field chambers (44.4 x 22.9 x 30.5 cm; Med Associates Inc.; St. Albans, VT) were individually located within sound-attenuating cubicles equipped with an exhaust fan that provided both ventilation and masking of external sounds. Time spent and distance travelled on each side of the chamber was measured with 4 parallel beams across the chamber floor.

### 2.3. Acoustic Startle Response (ASR)

On PND 29, approximately 10 min following the open field test, the acoustic startle response was assessed using a startle response system (S-R Lab; San Diego Instruments, San Diego, CA). Rats were placed in a cylindrical Plexiglas animal enclosure ((8” (L) x 3.5” (ID)) located within a sound-attenuating test chamber that included an exhaust fan, a sound source, and an internal light that was turned off during the test. At the start of each test, rats underwent a 5-min habituation period during which 60 dB of background white noise was present. The background noise was present during the entire test session. The test session consisted of 30 trials of a 250ms burst of a series of three randomized tones (100, 110, 120 dB) for 30 trials with a 30 s inter-trial interval. Startle amplitude was measured with a high-accuracy accelerometer mounted under the animal enclosure and was analyzed with SR-Lab software.

### 2.4. Light*/Dark Test*

On PND 30, animals underwent testing for anxiety-like behavior as measured by approach/avoidance behavior in the light/dark chamber. A dark box insert (44.4 x 22.9 x 30.5 cm) was placed in the left side of an open field chamber (23.31 x 27.31 x 20.32 cm; Med Associates Inc.; St. Albans, VT) to divide the chamber into a dark and light side (150 lux in center of light side). The chamber was located within sound attenuating cubicles equipped with an exhaust fan that provided both ventilation and masking of external sounds. Time spent and distance traveled on each side of the chamber was measured with 4 parallel beams across the chamber floor. Animals were habituated to the testing room for at least 20 min prior to the start of the 10 min test and each rat was placed in the light side facing the posterior wall.

### 2.5. Two-Bottle Choice Intermittent Access to Alcohol Drinking

On PND 31, rats began the two-bottle choice home cage alcohol drinking experiment. Rats were double housed by sex x litter x treatment and had one bottle of 20% ethanol and one bottle of water every other day for three weeks (9 sessions). Placement of bottles on the cage were switched for each session to avoid side preferences. Alcohol intake (g/kg) and water intake (mL) were calculated for each session.

### 2.6. Operant Alcohol Self-Administration Training

After completion of the home cage drinking phase, rats were undisturbed until approximately postnatal day 70 (young adulthood) when operant alcohol self-administration training began. Self-administration chambers (31 x 32 x 24 cm; Med Associates Inc.; St. Albans, VT) were individually located within sound-attenuating cubicles equipped with an exhaust fan that provided both ventilation and masking of external sounds. Chambers were fitted with a retractable lever on the left and right walls. A white cue light was centered 7-cm above each lever and a liquid receptacle was centered on each wall. Lever responses on the left lever (i.e. active/alcohol lever) activated a syringe pump (Med Associates) that delivered 0.1 ml of solution into the receptacle during a 1.66-s period in which both the white cue light and tone located above the active lever were activated. Responses on the right lever (i.e. inactive lever) had no programmed consequence. The chambers also had infrared photobeams, which divided the floor of the chamber into 4 zones to record general locomotor activity throughout each session. Rats (approximately PND 70, young adulthood) were trained on operant alcohol self-administration training. For training sessions, rats received two overnight self-administration training sessions with 20% alcohol reinforcer. During the first overnight (16 hr) session, rats were required to reach a criterion of 150 reinforcers and on the second overnight (16 hr) session rats were required to reach a criterion of 100 reinforcers (both nights were on a fixed ratio (FR-2) schedule).

#### Operant Alcohol Self-Administration

The timeline of the self-administration sessions is illustrated in Figure 1. Self-administration sessions (30 minutes, FR-2 schedule) took place 5 days per week (M-F) for 20 sessions (20% v/v, alcohol). Rats then rats underwent another phase of 4 sessions with a sucrose only reinforcer (1% w/v, sucrose), and then 7 sessions with a sweetened alcohol reinforcer (20% v/v, alcohol; 1% w/v, sucrose). Following the last session, rats underwent an abstinence period from alcohol self-administration for 2 weeks (14 d) in which rats remained in the home cage and no self-administration sessions were conducted. Rats resumed operant self-administration for 4 days (20% v/v, alcohol; 1% v/v, sucrose) to examine the reinitiation of alcohol self-administration. For this phase, rats were returned to alcohol self-administration for 4 days with the sweetened reinforcer (20% v/v, alcohol; 1% w/v, sucrose).

##### 2.6.1. Data Analysis

For basal corticosterone levels, distance traveled during open field test, % time in light side during light/dark test, total alcohol intake for two-bottle choice and alcohol self-administration, two-way ANOVAs were used to analyze main effects and interactions between prenatal alcohol exposure (PAE) and prenatal cannabinoid exposure (PCE) groups. For ASR, alcohol intake (g/kg) in the two-bottle choice sessions, and alcohol lever responses, intake and inactive lever responses across self-administration sessions, three-way RM ANOVAs were used to analyze main effects and interactions. Two-way ANOVAs were used to analyze interactions between PAE or PCE x session (or decibel for ASR). One-way ANOVAs were used to analyze individual main effects of PAE or PCE. During OFT, one female was excluded from analysis due to significant outlier.

## 3. Results

### The effects of gestational drug exposure on baseline corticosterone levels

#### Males

A two-way ANOVA showed a significant main effect of prenatal alcohol exposure (PAE) (Fig 2A, F(1,33) = 6.37, *p* < 0.05), with prenatal alcohol-exposed (PAE) adolescent males having significantly lower basal CORT levels as compared to prenatal water-exposed rats. There was no significant main effect of prenatal cannabinoid exposure (PCE) nor an PAE x PCE interaction, *p* > 0.05.

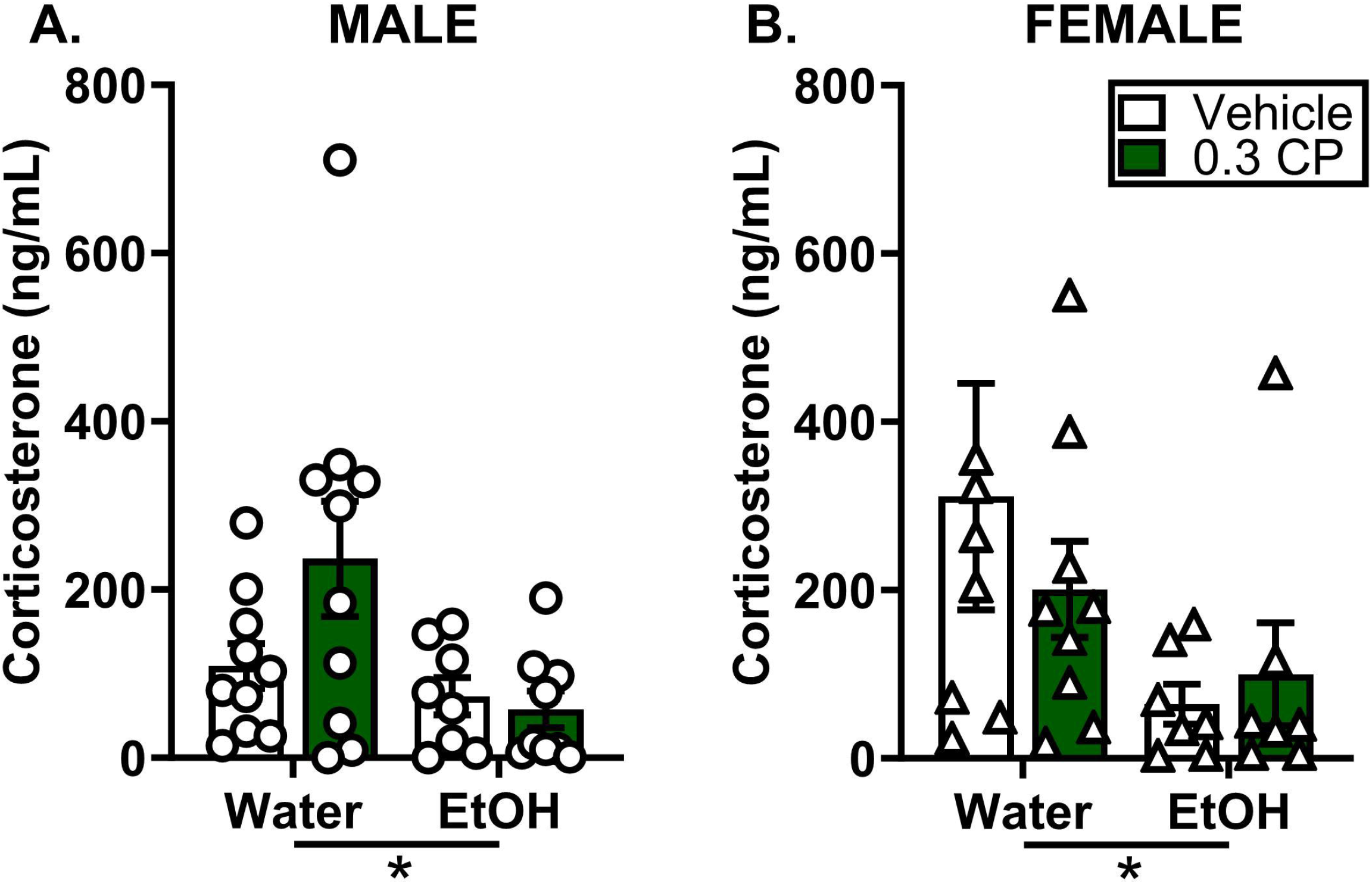

#### Females

Similarly in females, a two-way ANOVA showed a significant main effect of PAE (Fig 2B, F(1,27) = 4.36, *p* < 0.05), such that adolescent females exposed to prenatal alcohol had decreased basal CORT levels compared to females exposed to prenatal water. There was no significant effect of PCE or PAE x PCE interaction (Fig 2B, *p* > 0.05).

### The effects of gestational drug exposure on locomotor, acoustic startle response and anxiety-like behavior

#### Open field Test

##### Males

There was a significant main effect of PAE (Fig. 3A, F (1, 34) = 4.66, *p* < 0.05) showing greater distance traveled, which was driven by a PAE x PCE interaction (Fig 3A, F(1, 34) = 5.60, *p* < 0.05), showing that rats treated with PAE+PCE traveled more than the PCE-only group.

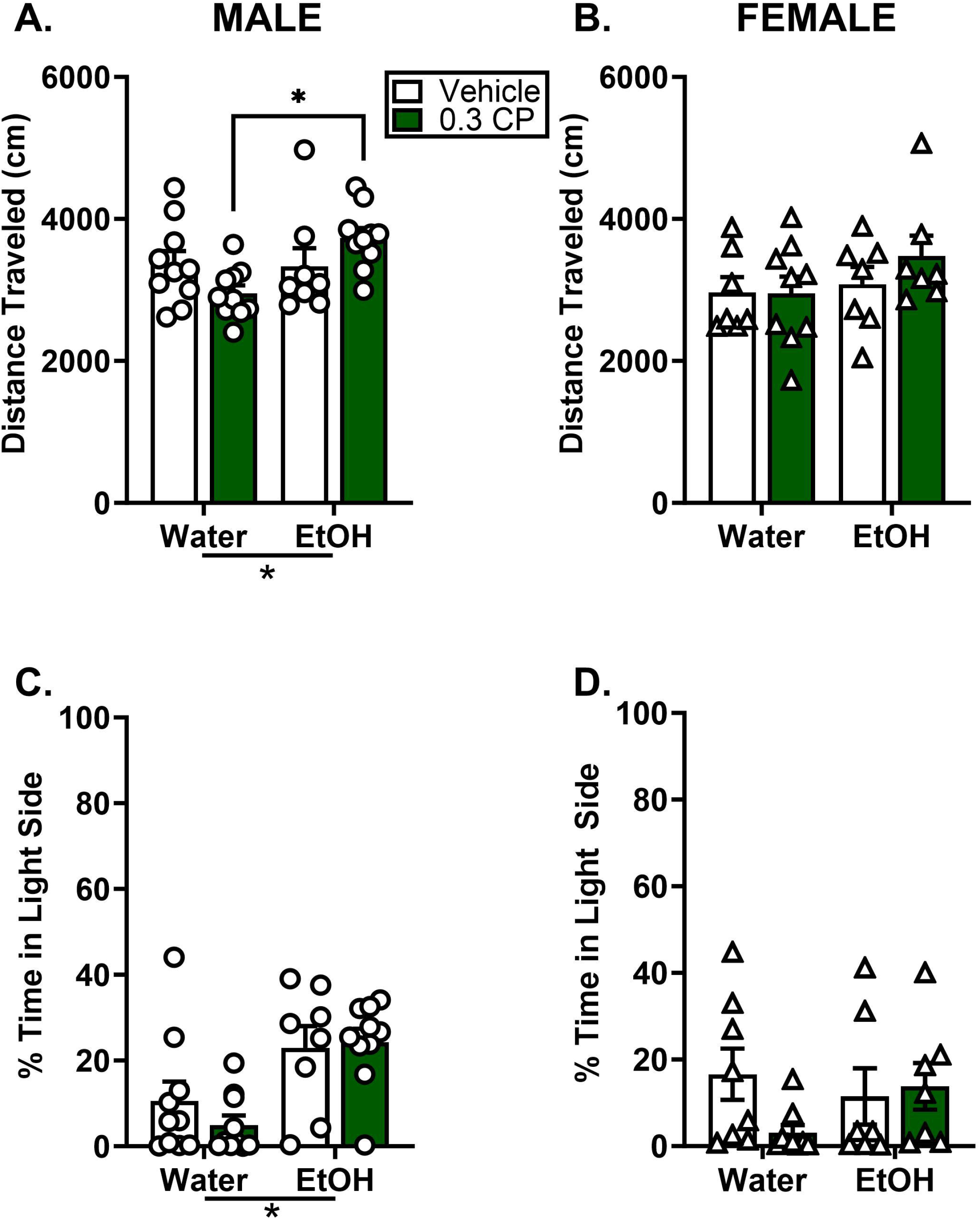

##### Females

Gestational drug exposure had no effect on locomotor behavior in the open field (Fig 3B, *p* > 0.05).

#### Acoustic Startle Response *(see Table 1)*

##### Males

A three-way RM ANOVA showed a significant main effect of decibel tones (Table 1, F (2, 68) = 111.7, *p* < 0.05) with increasing startle as the decibels increased as would be expected. There was a session x PAE x PCE interaction (Table 1, F (2,68) = 3.34, *p* < 0.05) with post hoc comparisons showing male rats exposed to PAE had a decrease startle response compared to controls at the 120dB level. There was also a PAE x PCE interaction (Table 1, F(1, 34) = 6.51, *p* < 0.05). No other main effects were observed.

##### Females

A three-way RM ANOVA showed a significant main effect of decibel tones (Table 1, F (2, 54) = 96.58, p < 0.05) with increasing startle as the decibels increased as would be expected. There was a significant PAE x PCE x decibel tone interaction (Table 1, F (2, 54) = 3.19, *p* < 0.05). There were no significant main effects of PAE, PCE or PAE x PCE x decibel interaction (p < 0.05). Two-way RM ANOVAs were used to follow up on the decibel x PAE x PCE interaction, to compare PAE and PCE startle amplitudes at each decibel separately. However, no differences were observed (*p* < 0.05).

#### Light/Dark Test

##### Males

Two-way ANOVA showed a significant main effect of PAE (Fig 3C, F(1, 34) = 18.46, *p* < 0.05) on % time spent in light side, such that males with PAE spent significantly more time in the light side compared to males exposed to gestational water. There was no significant effect of PCE or PAE x PCE interaction (*p* > 0.05).

##### Females

Gestational drug exposure had no effect on % time in the light side (Fig 3D, *p* > 0.05).

### Gestational drug exposure produces differences in alcohol intake (two-bottle choice) during adolescence

#### Males

The three-way ANOVA analysis of alcohol intake (g/kg) across sessions (3-d avg) showed a significant main effect of session (Fig 4A, F (2, 68) = 27.19, *p* < 0.05), with a decrease in alcohol consumption across the session. There was a significant main effect of PCE (Fig 4A, F (1, 34) = 5.03, *p* < 0.05), showing that PCE-treated rats consumed more alcohol. There were no main effect of PAE or significant interactions (*p* > 0.05). Analysis of total alcohol intake (g/kg) via two-way ANOVA showed a significant main effect of PCE (Fig 4B, F(1, 34) = 16.58, *p* < 0.05), such that PCE-males consumed a greater amount of alcohol compared to males exposed to vehicle. There were no significant differences in water intake (mL) between groups (Water + Veh: 173.15 ± 8.31 ml; Water + CP: 160.85 ± 7.13 ml; EtOH + Veh: 183.93 ± 11.12 ml; EtOH + CP: 170.06 ± 7.97 ml; *p* > 0.05).

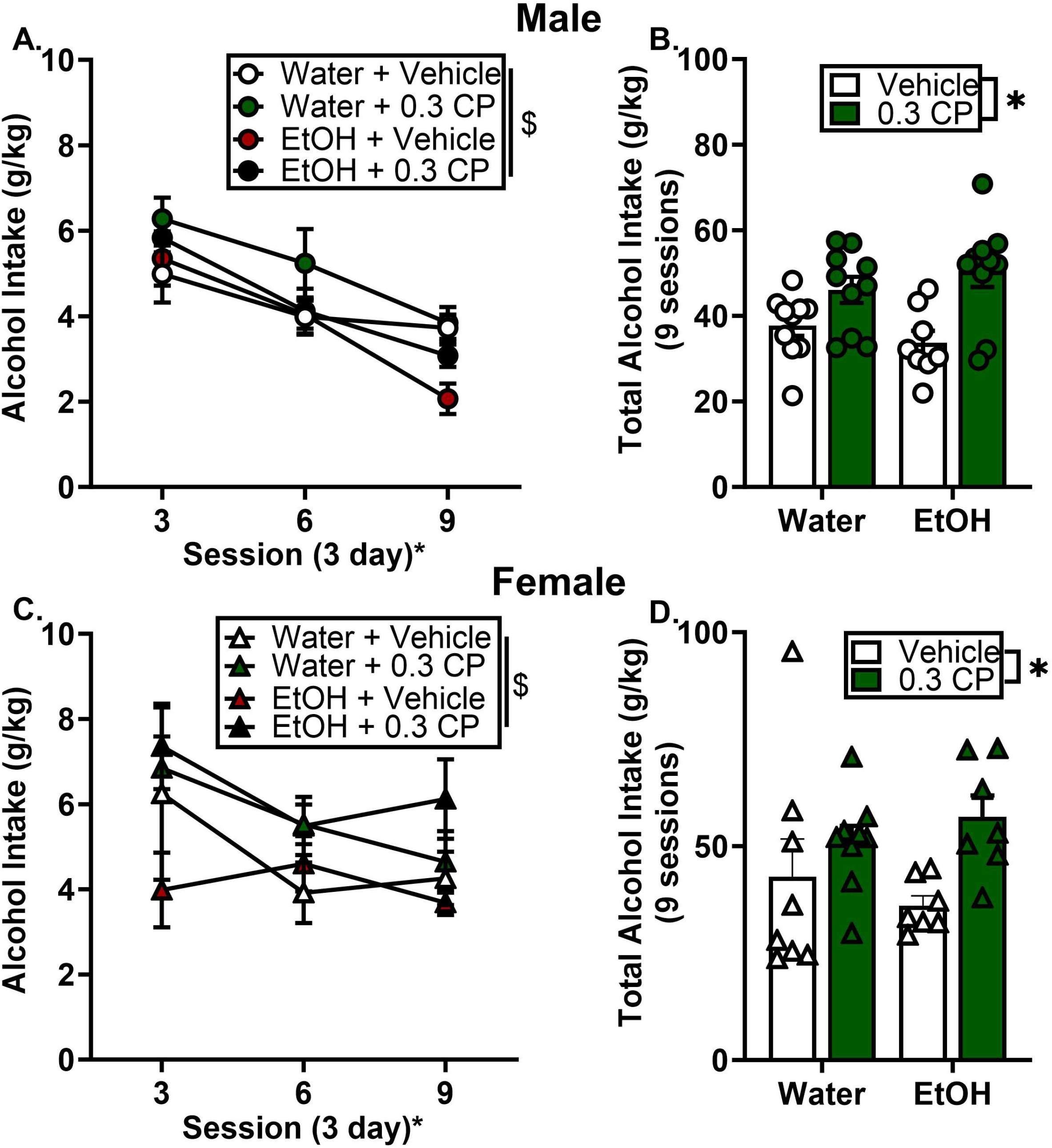

#### Females

Analysis of alcohol intake (g/kg) across sessions (3-d avg) showed a significant main effect of session (Fig 4C, F(2, 54) = 3.49, *p* < 0.05), with a general decrease in consumption across the sessions. There was a main effect of PCE (F(1, 27) = 6.18 *p* < 0.05), showing that female PCE rats showed increased alcohol intake during adolescence. There was no main effect of PAE or significant interactions (*p* > 0.05). Similar to males, analysis of total alcohol intake (g/kg) showed a significant main effect of PCE (Fig 4D, F(1, 27) = 6.58, *p* < 0.05), such that female PCE rats showed increased alcohol intake during adolescence. Interestingly, there was a significant PAE x PCE interaction for water intake (mL), showing that rats exposed to gestational alcohol + CP consumed more water compared to water + CP females (F (1, 27) = 7.26, *p* < .05; Water + Veh: 141.06 ± 4.38 ml; Water + CP: 118.94 ± 6.33 ml; EtOH + Veh: 132.40 ± 2.38 ml; EtOH + CP: 143.07 ± 9.06 ml).

### The effects of gestational drug exposure on alcohol intake during operant self-administration in adult females but not males

#### Males

During alcohol self-administration in adulthood, male rats showed a significant increase in self-administration across the sessions as confirmed by a significant main effect of session for alcohol lever responses (Fig. 5A, F (9, 306) = 30.52, *p* < 0.05) in the three-way ANOVA. There was also a significant PCE x session interaction (Fig. 5B, F (9, 306) = 2.13, *p* < 0.05), showing that male PCE rats had a general decrease in alcohol lever responses during alcohol self-administration; however this was not shown in total alcohol intake (g/kg). Two-way RM ANOVAs were used to follow up on the PCE x session interaction; however, post hoc comparisons tests did not show any differences between PCE groups across sessions (*p* > 0.05). Male rats showed no significant main effect of PAE or PCE, PAE x session, PAE x PCE or PAE x PCE x session interactions (Fig. 5B, *p* > 0.05). Alcohol intake g/kg (2-d avg) and inactive lever responses (2-d avg) are reported in Table 2.

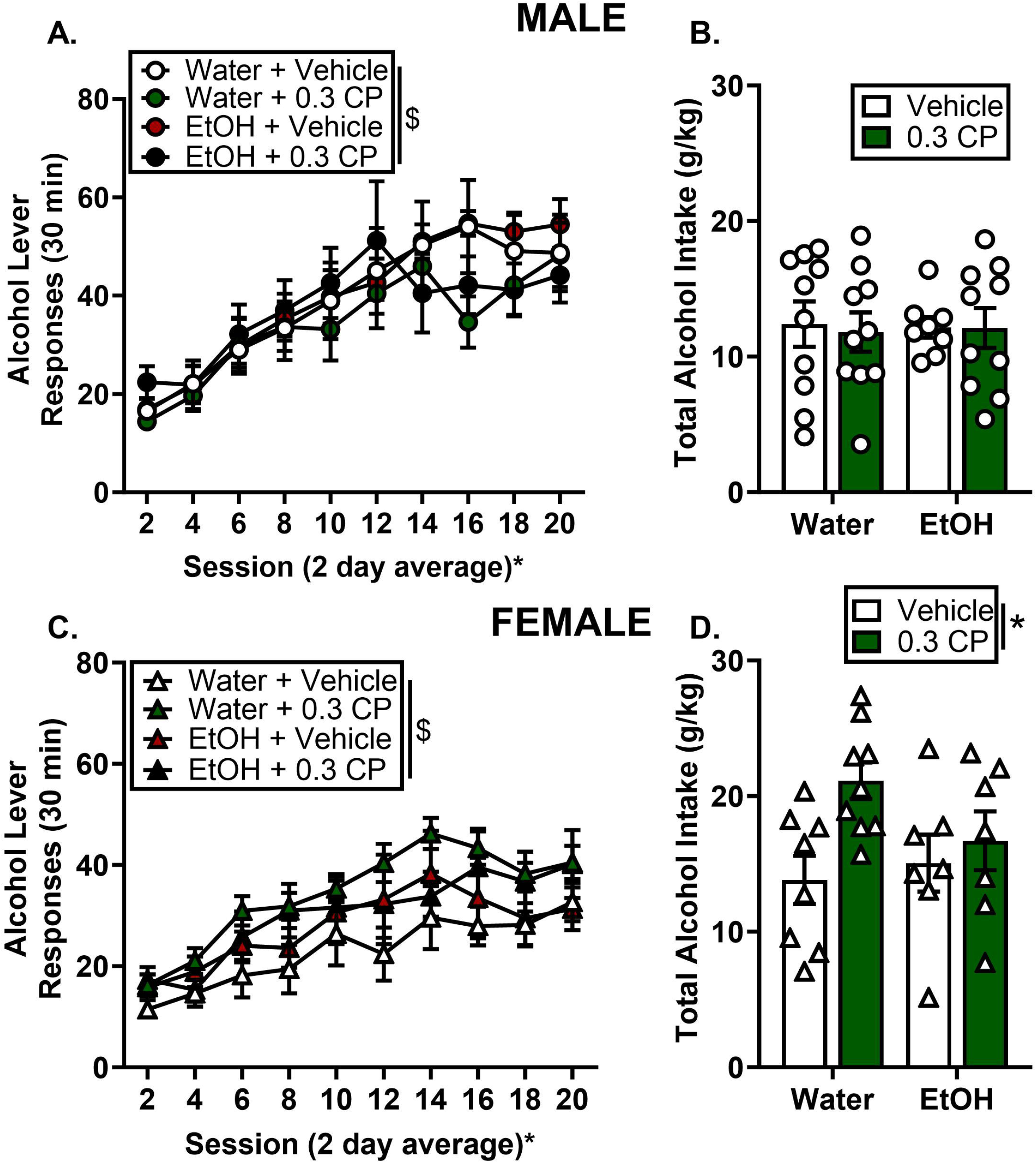

**Table 2.** Alcohol intake (g/kg) and ll11active lever reS1ponses during alcohol self admi11is1ration acquisition phase in males.

|  |  | Alcohol Intake (g/kg) |  |  |  |  |  |  |  |  |  |
| --- | --- | --- | --- | --- | --- | --- | --- | --- | --- | --- | --- |
| Group |  | 2 | 4 | 6 | 8 | 10 | 12 | 14 | 16 | 18 | 20 |
| Male | Water, Veh | 0.31±0.03 | 0.39±0.06 | 0.50±0.06 | 0.57±0.08 | 0.64±0.12 | 0.72±0.13 | 0.77±0.12 | 0.85±0.14 | 0.74±0.11 | 0.72±0.10 |
|  | Water, 0.3 CP | 0.28±0.04 | 0.34±0.06 | 0.55±0.11 | 0.61±0.13 | 0.58±0.12 | 0.73±0.12 | 0.74±0.08 | 0.60±0.08 | 0.74±0.10 | 0.75±0.09 |
|  | EtOH, Veh | 0.28±0.03 | 0.37±0.08 | 0.49±0.09 | 0.56±0.06 | 0.63±0.09 | 0.65±0.05 | 0.75±0.05 | 0.81±0.04 | 0.77±0.06 | 0.79±0.08 |
|  | EtOH, 0.3 CP | 0.39±0.05 | 0.37±0.06 | 0.53±0.09 | 0.61±0.10 | 0.70±0.12 | 0.82±0.17 | 0.65±0.11 | 0.67±0.08 | 0.64±0.08 | 0.67±0.07 |
| Inactive Lever Responses |  |  |  |  |  |  |  |  |  |  |  |
| Male | Water, Veh | 2.25±0.61 | 2.90±0.55 | 3.15±0.72 | 3.95±1.14 | 2.85±1.15 | 1.25±0.27 | 2.50±1.02 | 3.20±0.93 | 0.90±0.26 | 1.80±0.50 |
|  | Water, 0.3 CP | 2.50±0.54 | 1.90±0.62 | 2.75±0.79 | 3.00±0.91 | 1.10±0.45 | 0.95±0.24 | 4.55±0.58 | 1.35±0.55 | 0.55±0.27 | 1.1±0.40 |
|  | EtOH, Veh | 2.25±0.62 | 3.25±0.53 | 3.63±0.62 | 2.69±0.61 | 3.25±0.60 | 2.50±0.63 | 2.06±0.57 | 2.31±0.35 | 1.81±0.63 | 1.81±0.46 |
|  | EtOH, 0.3 CP | 2.60±0.55 | 2.80±0.55 | 2.15±0.65 | 3.05±1.07 | 2.60±0.99 | 1.85±0.60 | 1.90±0.63 | 1.60±0.63 | 1.40±0.31 | 2.35±0.54 |

Mean ± SEM. \* main effect of session, ^ main effect of PAE, \$ main effect of PCE, & significant session x PAE interaction, # significant session x PCE interaction, + significant PAE x PCE interaction, † significant decible x PAE x PCE interaction, p < 0.05.

#### Females

The three-way ANOVA analysis of self-administration in females showed a significant main effect of session for alcohol lever responses (Fig. 5C, F (9, 243) = 19.10, *p* < 0.05) such that alcohol lever responses increased overtime for all exposure and treatment groups. There was also a main effect of PCE (Fig. 5C, F (1, 27) = 4.62, *p* < 0.05), such that across all sessions PCE females showed a general increase in self-administration compared to controls. Furthermore, examination of total alcohol intake (g/kg), showed a significant main effect of PCE (Fig 5D, F (1, 27) = 6.01, *p* < 0.05). Therefore, gestational CP-exposed females self-administered more total alcohol (g/kg) than did the vehicle-exposed females. There was no significant main effect of PAE or PAE x PCE interaction (*p* > 0.05). These results suggest gestational exposure to CP can lead to increases in alcohol intake in adulthood, specifically in females. Alcohol intake g/kg (2-d avg) and inactive lever responses (2-d avg) are reported in Table 3.

**Table 3.** Alcohol intake (g/kg) and ll11active lever reS1ponses during alcohol self admi11is1ration acquisition phase in females.

|  |  | Alcohol Intake (g/kg) |  |  |  |  |  |  |  |  |  |
| --- | --- | --- | --- | --- | --- | --- | --- | --- | --- | --- | --- |
| Group |  | 2 | 4 | 6 | 8 | 10 | 12 | 14 | 16 | 18 | 20 |
| Females | Water, Veh | 0.39±0.06 | 0.50±0.09 | 0.53±0.13 | 0.58±0.15 | 0.70±0.13 | 0.65±0.14 | 0.93±0.14 | 0.84±0.152 | 0.81±0.12 | 0.98±0.11 |
|  | Water, 0.3 CP | 0.55±0.09 | 0.65±0.07 | 0.99±0.10 | 1.06±0.10 | 1.05±0.10 | 1.21±0.13 | 1.34±0.15 | 1.32±0.10 | 1.16±0.12 | 1.22±0.14 |
|  | EtOH, Veh | 0.47±0.08 | 0.54±0.07 | 0.66±0.08 | 0.64±0.10 | 0.85±0.19 | 0.84±0.19 | 0.94±0.18 | 0.90±0.18 | 0.81±0.17 | 0.88±0.14 |
|  | EtOH, 0.3 CP | 0.50±0.04 | 0.44±0.07 | 0.75±0.14 | 0.86±0.14 | 0.90±0.14 | 0.92±0.17 | 0.90±0.14 | 1.08±0.17 | 0.95±0.14 | 1.05±0.13 |
|  |  | Inactive Lever Responses |  |  |  |  |  |  |  |  |  |
| Females | Water, Veh | 1.31±0.54 | 2.06±0.83 | 1.63±0.65 | 3.56±2.05 | 2.63±1.03 | 3.88±2.88 | 1.75±0.86 | 2.69±1.68 | 1.75±1.19 | 1.19±0.71 |
|  | Water, 0.3 CP | 2.00±0.63 | 1.67±0.34 | 2.17±0.52 | 1.67±0.40 | 2.61±0.79 | 1.67±0.40 | 4.72±1.49 | 1.67±0.51 | 1.83±0.64 | 1.61±0.39 |
|  | EtOH, Veh | 2.00±0.44 | 3.86±1.5 | 2.07±0.69 | 4.29±1.22 | 3.36±0.84 | 2.93±1.03 | 3.36±1.35 | 1.79±0.53 | 1.36±0.56 | 2.71±0.78 |
|  | EtOH, 0.3 CP | 3.21±0.84 | 3.00±0.85 | 3.36±1.07 | 2.07±0.70 | 4.29±1.16 | 3.21±1.31 | 3.36±1.05 | 4.00±1.50 | 1.93±0.80 | 3.57±1.33 |

Mean ± SEM. \* main effect of session, ^ main effect of PAE, \$ main effect of PCE, & significant session x PAE interaction, # significant session x PCE interaction, + significant PAE x PCE interaction, † significant decible x PAE x PCE interaction, p < 0.05.

### Change in reinforcer produces altered sensitivity in males but not females

#### Males

To examine how a change in reinforcer might alter responding during self-administration, rats were introduced to a non-alcohol 1% sucrose reinforcer after the final 20% alcohol self-administration session. First, a two-way ANOVA comparing the average of the last two alcohol sessions showed no main effects of PAE, PCE or PAE x PCE interactions for alcohol lever responses (Fig. 6A: 2d avg, last two 20% alcohol sessions, *p* > 0.05), meaning that all exposure groups had similar levels of alcohol self-administration before the change in reinforcer. During the 1% sucrose sessions there was a significant PAE x PCE interaction (Fig. 6B, F (1, 34) = 5.84, *p* < 0.05), session x PCE interaction (Fig. 6B, F (3, 102) = 3.13, *p* < 0.05) and a main effect of PCE (Fig. 6B, F (1,34) = 5.47, *p* < 0.05) and session (Fig. 6B, F (3,102) = 2.82, *p* < 0.05). Multiple comparisons tests showed male rats exposed to PCE showed reduced lever responses during the last 3 days out of the 4 days on 1% sucrose compared to controls (Figure 6B, *p* < 0.05). There was no main effect of PAE, PAE x session interaction, or PAE x PCE x session interaction (*p* > 0.05). Next, 20% alcohol was added to the 1% sucrose. There was a PCE x session interaction (Fig. 6B, F (6, 204) = 2.32, *p* < 0.05), such that during 20% alcohol, 1% sucrose self-administration days, male rats exposed to gestational CP continued to make fewer alcohol lever responses than did the control rats; however multiple comparisons tests did not show significant differences between PCE and controls during each 20%, 1% session (*p* > 0.05). There was no main effect of PAE, nor a PAE x PCE interaction for alcohol lever responses (Fig. 6C, *p* > 0.05).

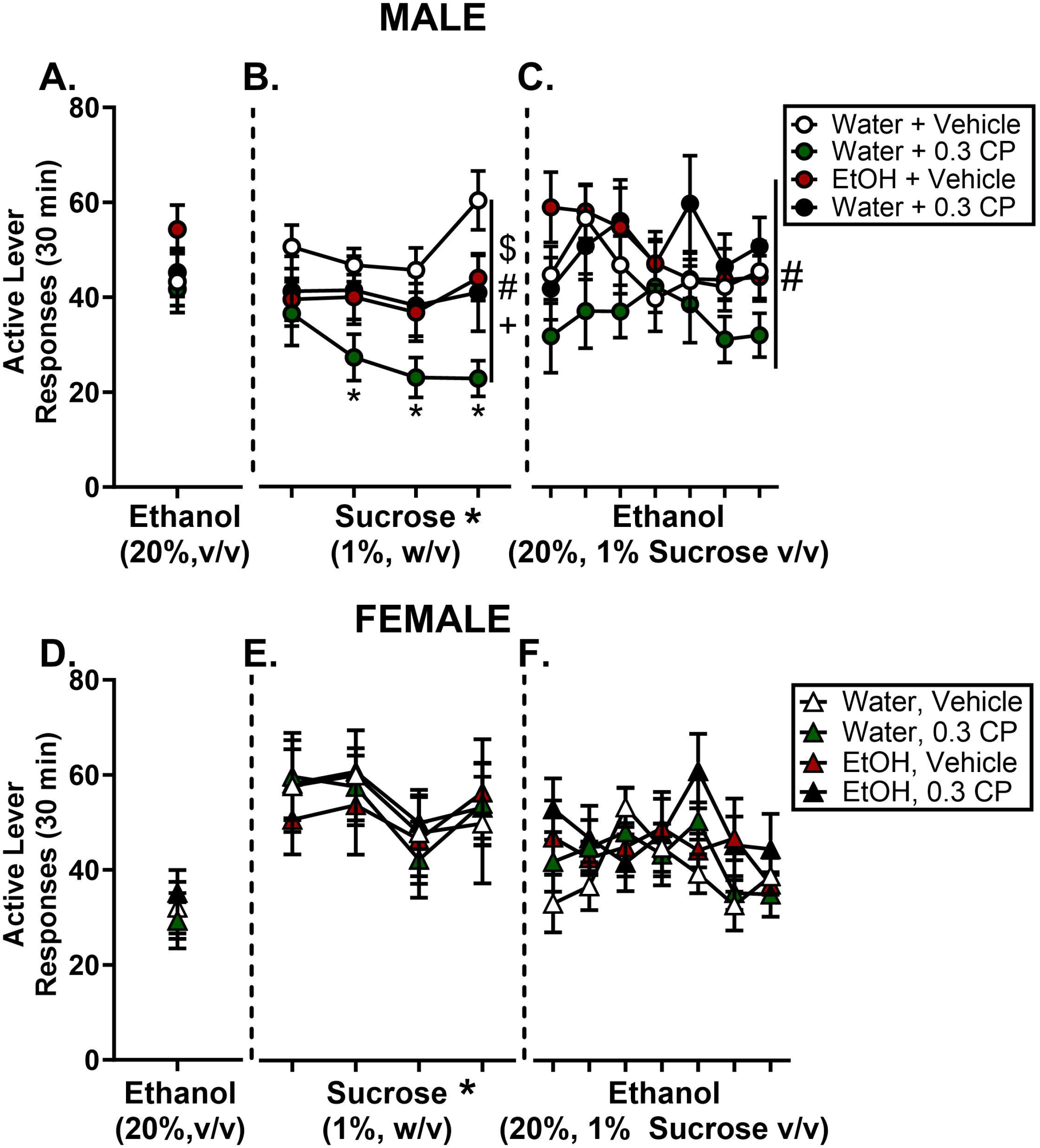

#### Females

There were no significant main effects of PCE or PAE or interactions during all three reinforcer phases of self-administration (Fig. 6D, 2d avg, last two 20% alcohol sessions; Fig. 6E, 1% sucrose; Fig. 7F, 20% alcohol + 1% sucrose, *p* > 0.05). During the 1% sucrose self-administration days, there was a significant main effect of session (Fig. 6E, F (3, 84) = 3.50, *p* < 0.05), such that alcohol lever responses decreased over 1% sucrose sessions.

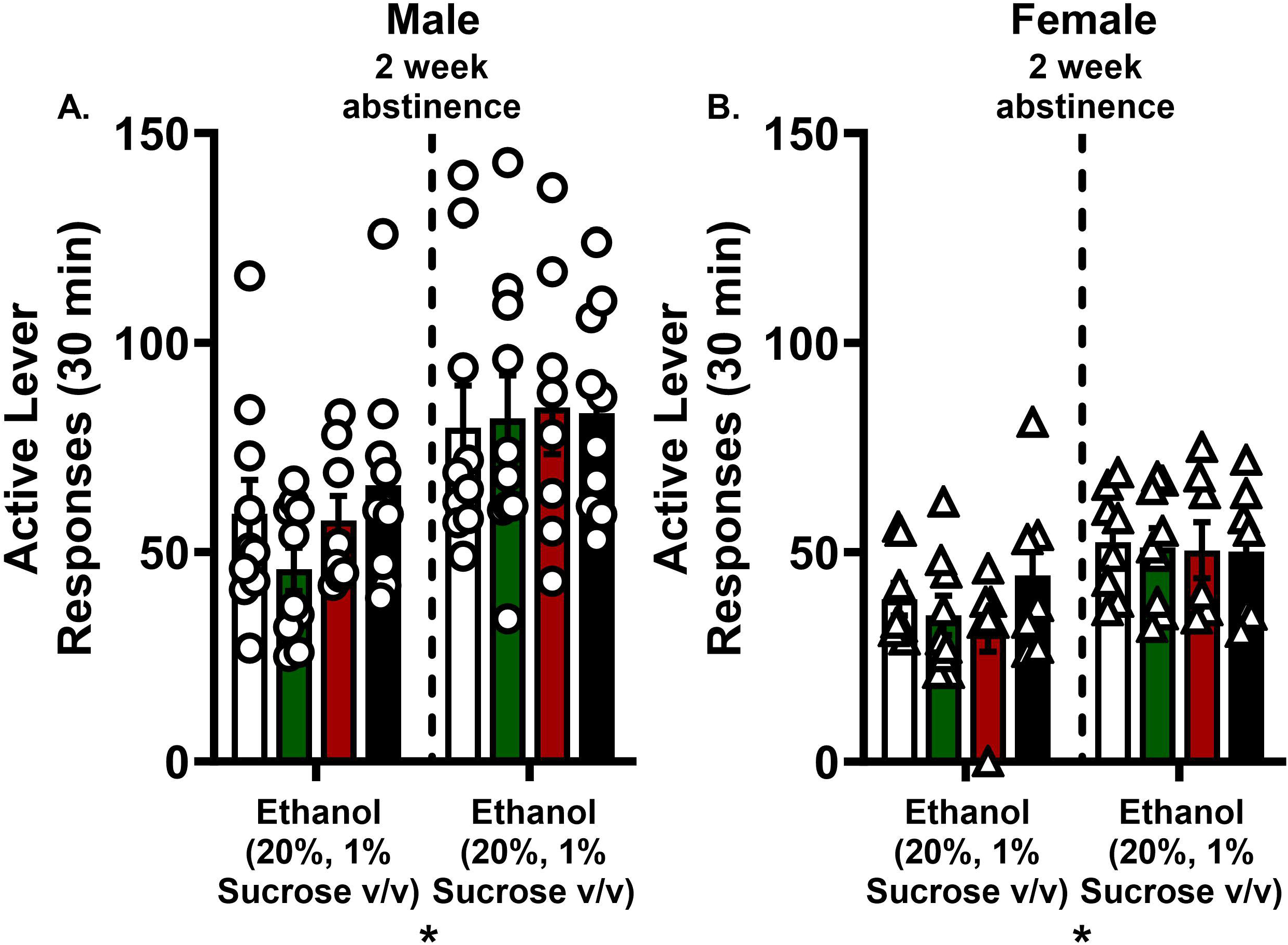

### Reinitiation of self-administration after abstinence period

#### Males

To examine reinitiation of alcohol drinking after the abstinence period, we examined alcohol lever responses before and after a 2-week abstinence period from self-administration sessions. There was a main effect of session (Fig. 7A, F (1, 34) = 26.63, *p* < 0.05), such that males regardless of gestational drug exposure showed an increase in alcohol lever responses after the 2-week abstinence period. However, there were no significant main effects of PAE, PCE, nor interactions (*p* > 0.05).

#### Females

Similar to males, there was also a main effect of session (Fig. 7B, F (1, 27) = 20.38, *p* < 0.05, such that females regardless of gestational drug exposure showed an increase in alcohol lever responses after the 2-week abstinence period. However, there were no significant main effects of PAE, PCE, nor interactions (*p* > 0.05).

## 4. Discussion

In the current study, we sought to determine the consequences of prenatal alcohol + SCB co-exposure on neuroendocrine and behavioral adaptations in male and female adolescent and adult rats and whether these effects were additive or synergistic. Overall, the results show persistent effects of either prenatal alcohol exposure (PAE) or prenatal cannabinoid exposure (PCE) that varied by sex, but no synergistic effects were observed. Specifically, PAE induced an alteration in basal corticosterone and arousal in both males and females; however, only males were sensitive to behavioral disruptions in locomotion and anxiety-like behavior. In contrast, female rats exposed to PCE showed increased alcohol drinking during adolescence and adulthood. Lastly, male PCE rats showed reduced sensitivity to sucrose reinforcement, potentially indicative of an anhedonic response. Overall, while PAE and PCE produced varying behavioral effects across adolescence and adulthood, no synergistic effects were observed.

Both alcohol and cannabinoid exposure alone during gestation can produce enduring neurobiological changes, especially of the hypothalamic-pituitary-adrenal (HPA) axis (Biggio et al., 2018; del Arco et al., 2000; Franks, Berry, & DeFranco, 2020; Wieczorek, Fish, O’Leary-Moore, Parnell, & Sulik, 2015), which plays a major part in modulating neuroendocrine functioning. Interestingly, we found that both adolescent male and female rats treated with PAE, but not PCE, had lower basal corticosterone levels. These results are consistent with other prenatal alcohol exposure studies (Hellemans, Sliwowska, Verma, & Weinberg, 2010; Wieczorek et al., 2015), showing dysregulated HPA axis function in adolescent offspring. Exposure to a single binge-like level of alcohol during gastrulation induces dysmorphologies in ventral midline regions, including the pituitary and hypothalamus (Boschen, Fish, & Parnell, 2021; Godin, Dehart, Parnell, O’Leary-Moore, & Sulik, 2011), which may contribute to HPA axis dysregulation. In some species, acute exposure to SCBs also induces midline brain abnormalities (Gilbert et al., 2016), however no effects on basal corticosterone levels were observed in the present study. Therefore, it will be important to follow up with an examination of HPA axis function in response to a challenge, such as a stressor, to get a more definitive understanding of the consequences of prenatal SCB exposure.

Children and adolescents with prenatal alcohol or cannabinoid exposure display anxiety-like behavior and hyperarousal (Jutras-Aswad, DiNieri, Harkany, & Hurd, 2009; Mulligan & Hamre, 2023; Nashed, Hardy, & Laviolette, 2021; Weile et al., 2020; Weiss, Jonn_-_Seed, & Harris_-_Muchell, 2007). In the current study, males treated with PAE + PCE showed an increase in locomotor behavior in the open field test relative to males exposed to PCE alone; however there was no evidence of hyperactivity when compared to the vehicle control group. While this finding supports an interaction between prenatal alcohol and cannabinoid exposure it is not indicative of a hyperactivity profile. Recent studies on combined alcohol and cannabinoid exposure found the male rats born to dams exposed to THC e-cigarette and vaporized alcohol throughout pregnancy (GD5-20) were hyperactive in the open field test (Breit, Rodriguez, Lei, Hussain, & Thomas, 2022). However, in a model for late gestational exposure (PD 4-9, third trimester equivalent) alcohol caused hyperactivity in adolescent rats, but there was no additive effect of alcohol and combined with CP-55,940 (Breit, Zamudio, & Thomas, 2019). Together, these findings suggest that while there may be lasting consequences of PAE + PCE on locomotor behavior, this likely depends on the timing of exposure.

In relation to anxiety-like behavior, male rats exposed to PAE showed an increase in the percent time spent in the light side of a compartment during the light/dark test, suggesting decreased anxiety-like behavior. Given that male rats exposed to PAE also showed reduced basal CORT levels, it is possible that increased time spent in the light side may be related to an altered stress/anxiety response or behavioral disinhibition. In relation to arousal, gestational drug exposure had minimal effects on startle response in both sexes, with the exception that males in the PAE group showed decreased startle response compared to the vehicle control group at the highest decibel tone (120 dB), suggesting hyperarousal. A decrease in arousal state could be indicative of a maladaptation involved in brain regions that regulate arousal or excitation in response to stimuli.

Following the behavioral screens, we sought to determine whether PAE or PCE influences alcohol drinking-related behavior during adolescence. As prenatal alcohol and cannabinoid exposure alone induces long lasting neuronal changes in frontocortical and limbic system brain regions (Bellinger, Davidson, Bedi, & Wilce, 2002; Gilbert et al., 2016; Rice et al., 2012; S. K. Rouzer, Cole, Johnson, Varlinskaya, & Diaz, 2017) that are involved in regulating addiction-related neural circuitry, we hypothesized that PAE + PCE would produce synergistic effects, resulting in increased alcohol intake during adolescence. In contrast to the adolescent behavioral screens, we found that during two-bottle choice, we found that in both males and females treated with PCE had greater total alcohol intake. We also saw that females exposed to PAE + PCE consumed greater water intake compared to females treated with water + CP. These results show that prenatal treatment of synthetic cannabinoids regardless of co-exposure to alcohol are enough to produce increases in adolescent alcohol drinking, which is an important finding for the growing body of research investigating long-term consequences of prenatal cannabinoid exposure.

An important feature of this study was that drinking behavior was also examined in adulthood, using an operant self-administration procedure. Both male and female rats underwent alcohol self-administration for approximately 4 weeks (total of 20 sessions). Males exposed to PCE showed a slight reduction in alcohol lever responses; however, total alcohol intake was not altered. In contrast to the males, PCE females showed increased alcohol self-administration and had a greater total alcohol intake. This finding is consistent with the two-bottle choice data in which the same group showed increased alcohol intake. Therefore, we find consistent and lasting effects of PCE exposure in alcohol consumption in female rats that persist from adolescence into adulthood. While we expected combination PAE + PCE to increase alcohol drinking in adolescence and adulthood, this was not the observed outcome. It is known that the endocannabinoid system regulates dopamine (DA) neuronal activity within the VTA (Melis et al., 2014; Serra, Aroni, Bortolato, Frau, & Melis, 2023), specifically through disinhibition of GABAergic neurons located in the rostromedial tegmental nucleus that regulate DA cells in the VTA (Lecca et al., 2011). Importantly, if endocannabinoids produce disinhibition of GABAergic neurons leading to greater activity of DA neurons in the VTA, this suggests a potential explanation on how endocannabinoids may contribute to enhanced DA cell activity and increased drug-seeking behavior; or increased alcohol drinking.

In order to determine sensitivity to a non-drug reinforcer, after the 20 sessions of alcohol (20%, v/v) self-administration, a sucrose (1%, w/v) reinforcer was introduced for one week. Males and females responded differently to this new reinforcer; males continued to respond for sucrose reinforcement at the same rate as alcohol reinforcement, while the females nearly doubled their response rates and maintained higher response rates when alcohol was reintroduced. In females, there was no effect of PAE or PCE on responding during the sucrose reinforcement phase of the experiment. In males, across the 4 sessions rats from each of the prenatal exposure groups tended to make fewer responses for sucrose reinforcement than did the control rats, although this suppressive effect was largest in the CP-alone group. This data pattern suggests that male rats exposed to PCE in particular, are less sensitive to the reinforcing effects of sucrose. Further, when alcohol was introduced to the sucrose solution the same rats exposed to PCE continued to make fewer responses throughout the duration of the self-administration sessions. This finding suggests that in males, PCE, regardless of co-exposure to alcohol, may blunt the stimulus value of sucrose and sweetened alcohol. On the basis of the sucrose responding data in males, it is possible that PCE affected ventral midline structures (Godin et al., 2011), such as the nucleus accumbens, which plays a major role in reward processing, but further studies are required to address this potential mechanism. Lastly, we investigated whether PCE and PAE would increase motivation to self-administer alcohol and alcohol-seeking behavior after an abstinence period. However, there was no effect of gestational drug exposure on abstinence-increased responding for sweetened alcohol. These data suggest that PCE-induced deficits in responding for sucrose can be overcome by increasing motivational drive through a period of abstinence.

It is important to discuss some limitations of these experiments. The current study examined a single dose of CP 55,940 (0.3 mg/kg) and alcohol (2.0 g/kg). Although these doses are within the range used by previous studies (0.1 to 0.4 mg/kg) (Breit et al., 2022; Breit et al., 2019), follow up experiments testing lower and higher dose combinations of SCB and alcohol will be important to address a broader spectrum of potential behavioral outcomes. Follow up studies may consider examining persistence of alcohol self-administration under various conditions such quinine adulteration. Also, inclusion of a sucrose self-administration control group would be beneficial to examine if increases in alcohol self-administration in response to PCE exposure is specific to alcohol, and whether the decrease in sucrose self-administration observed emerges if there is no alcohol history.

It will be important for future work to begin to investigate the neurobiological changes associated with this drug exposure. Two of the most important findings in the current study were blunted basal CORT levels due to PAE exposure and increases in alcohol consumption in adolescence and adulthood, specifically in females. Therefore, follow up studies can focus on examining neuronal adaptations within the HPA axis, specifically to probe dysregulated neuroendocrine responses associated with gestational drug exposure. Also, ventral midline structures such as the mPFC, nucleus accumbens and ventral tegmental area will be an important to focus on as they play a major role in reward processing.

## Conclusion

These studies utilize a unique approach to investigate the effects of gestational drug exposure on behavior by targeting drug exposure during gastrulation and neurulation, two important developmental periods for the formation of the brain. The current study shows that gestational exposure to alcohol or a synthetic cannabinoid each produces distinct effects on the offspring. We found that alcohol affected adolescent behavior and basal corticosterone levels more than cannabinoid exposure, while cannabinoid exposure was more likely to affect measures of alcohol drinking. These results emphasize the potential for negative consequences of PCE and PAE as well as co-exposure PAE + PCE on lasting behavioral adaptations in adolescence and adulthood, which has been understudied.

## Data availability

Available on request.

## Acknowledgements

none

## Bibliography

Bellinger, F. P., Davidson, M. S., Bedi, K. S., & Wilce, P. A. (2002). Neonatal ethanol exposure reduces AMPA but not NMDA receptor levels in the rat neocortex. Brain Res Dev Brain Res, 136(1), 77–84. doi:10.1016/s0165-3806(02)00363-2

Biggio, F., Talani, G., Locci, V., Pisu, M. G., Boero, G., Ciarlo, B., Grayson, D. R., & Serra, M. (2018). Low doses of prenatal ethanol exposure and maternal separation alter HPA axis function and ethanol consumption in adult male rats. Neuropharmacology, 131, 271–281. doi:10.1016/j.neuropharm.2017.12.005

Boschen, K. E., Fish, E. W., & Parnell, S. E. (2021). Prenatal alcohol exposure disrupts Sonic hedgehog pathway and primary cilia genes in the mouse neural tube. Reproductive Toxicology, 105, 136–147.

Breit, K. R., Rodriguez, C. G., Lei, A., Hussain, S., & Thomas, J. D. (2022). Effects of prenatal alcohol and delta-9-tetrahydrocannabinol exposure via electronic cigarettes on motor development. Alcohol Clin Exp Res, 46(8), 1408–1422. doi:10.1111/acer.14892

Breit, K. R., Zamudio, B., & Thomas, J. D. (2019). Altered motor development following late gestational alcohol and cannabinoid exposure in rats. Neurotoxicol Teratol, 73, 31–41. doi:10.1016/j.ntt.2019.03.005

Chang, J. C., Tarr, J. A., Holland, C. L., De Genna, N. M., Richardson, G. A., Rodriguez, K. L., Sheeder, J., Kraemer, K. L., Day, N. L., & Rubio, D. (2019). Beliefs and attitudes regarding prenatal marijuana use: perspectives of pregnant women who report use. Drug and alcohol dependence, 196, 14–20.

Coleman-Cowger, V. H., Schauer, G. L., & Peters, E. N. (2017). Marijuana and tobacco co-use among a nationally representative sample of US pregnant and non-pregnant women: 2005–2014 National Survey on Drug Use and Health findings. Drug and alcohol dependence, 177, 130–135.

Cook, J. L. (2020). Effects of prenatal alcohol and cannabis exposure on neurodevelopmental and cognitive disabilities Handbook of Clinical Neurology (Vol. 173, pp. 391–400): Elsevier.

Day, N. L., Richardson, G. A., Goldschmidt, L., Robles, N., Taylor, P. M., Stoffer, D. S., Cornelius, M. D., & Geva, D. (1994). Effect of prenatal marijuana exposure on the cognitive development of offspring at age three. Neurotoxicol Teratol, 16(2), 169–175. doi:10.1016/0892-0362(94)90114-7

De Genna, N. M., Willford, J. A., & Richardson, G. A. (2022). Long-term effects of prenatal cannabis exposure: Pathways to adolescent and adult outcomes. Pharmacol Biochem Behav, 214, 173358. doi:10.1016/j.pbb.2022.173358

del Arco, I., Muñoz, R., Rodríguez De Fonseca, F., Escudero, L., Martín-Calderón, J. L., Navarro, M., & Villanúa, M. A. (2000). Maternal exposure to the synthetic cannabinoid HU-210: effects on the endocrine and immune systems of the adult male offspring. Neuroimmunomodulation, 7(1), 16–26. doi:10.1159/000026416

Elsohly, M. A., Gul, W., Wanas, A. S., & Radwan, M. M. (2014). Synthetic cannabinoids: analysis and metabolites. Life Sci, 97(1), 78–90. doi:10.1016/j.lfs.2013.12.212

England, L. J., Bennett, C., Denny, C. H., Honein, M. A., Gilboa, S. M., Kim, S. Y., Guy, G. P., Jr., Tran, E. L., Rose, C. E., Bohm, M. K., & Boyle, C. A. (2020). Alcohol Use and Co-Use of Other Substances Among Pregnant Females Aged 12-44 Years - United States, 2015-2018. MMWR Morb Mortal Wkly Rep, 69(31), 1009–1014. doi:10.15585/mmwr.mm6931a1

Fish, E. W., Murdaugh, L. B., Zhang, C., Boschen, K. E., Boa-Amponsem, O., Mendoza-Romero, H. N., Tarpley, M., Chdid, L., Mukhopadhyay, S., Cole, G. J., Williams, K. P., & Parnell, S. E. (2019). Cannabinoids Exacerbate Alcohol Teratogenesis by a CB1-Hedgehog Interaction. Sci Rep, 9(1), 16057. doi:10.1038/s41598-019-52336-w

Franks, A. L., Berry, K. J., & DeFranco, D. B. (2020). Prenatal drug exposure and neurodevelopmental programming of glucocorticoid signalling. J Neuroendocrinol, 32(1), e12786. doi:10.1111/jne.12786

Fried, P., & Smith, A. (2001). A literature review of the consequences of prenatal marihuana exposure: an emerging theme of a deficiency in aspects of executive function. Neurotoxicology and teratology, 23(1), 1–11.

Gilbert, M. T., Sulik, K. K., Fish, E. W., Baker, L. K., Dehart, D. B., & Parnell, S. E. (2016). Dose-dependent teratogenicity of the synthetic cannabinoid CP-55,940 in mice. Neurotoxicology and teratology, 58, 15–22.

Godin, E. A., Dehart, D. B., Parnell, S. E., O’Leary-Moore, S. K., & Sulik, K. K. (2011). Ventromedian forebrain dysgenesis follows early prenatal ethanol exposure in mice. Neurotoxicol Teratol, 33(2), 231–239. doi:10.1016/j.ntt.2010.11.001

Hellemans, K. G., Sliwowska, J. H., Verma, P., & Weinberg, J. (2010). Prenatal alcohol exposure: fetal programming and later life vulnerability to stress, depression and anxiety disorders. Neuroscience & Biobehavioral Reviews, 34(6), 791–807.

Huizink, A. (2014). Prenatal cannabis exposure and infant outcomes: overview of studies. Progress in Neuro-Psychopharmacology and Biological Psychiatry, 52, 45–52.

Jutras-Aswad, D., DiNieri, J. A., Harkany, T., & Hurd, Y. L. (2009). Neurobiological consequences of maternal cannabis on human fetal development and its neuropsychiatric outcome. European archives of psychiatry and clinical neuroscience, 259, 395–412.

Lecca, S., Melis, M., Luchicchi, A., Ennas, M. G., Castelli, M. P., Muntoni, A. L., & Pistis, M. (2011). Effects of Drugs of Abuse on Putative Rostromedial Tegmental Neurons, Inhibitory Afferents to Midbrain Dopamine Cells. Neuropsychopharmacology, 36(3), 589–602. doi:10.1038/npp.2010.190

Martínez-Peña, A. A., Perono, G. A., Gritis, S. A., Sharma, R., Selvakumar, S., Walker, O. S., Gurm, H., Holloway, A. C., & Raha, S. (2021). The Impact of Early Life Exposure to Cannabis: The Role of the Endocannabinoid System. Int J Mol Sci, 22(16). doi:10.3390/ijms22168576

Melis, M., Sagheddu, C., Felice, M. D., Casti, A., Madeddu, C., Spiga, S., Muntoni, A. L., Mackie, K., Marsicano, G., Colombo, G., Castelli, M. P., & Pistis, M. (2014). Enhanced Endocannabinoid-Mediated Modulation of Rostromedial Tegmental Nucleus Drive onto Dopamine Neurons in Sardinian Alcohol-Preferring Rats. The Journal of Neuroscience, 34(38), 12716–12724. doi:10.1523/jneurosci.1844-14.2014

Mereu, G., Fà, M., Ferraro, L., Cagiano, R., Antonelli, T., Tattoli, M., Ghiglieri, V., Tanganelli, S., Gessa, G. L., & Cuomo, V. (2003). Prenatal exposure to a cannabinoid agonist produces memory deficits linked to dysfunction in hippocampal long-term potentiation and glutamate release. Proceedings of the National Academy of Sciences, 100(8), 4915–4920.

Metrik, J., & Patel, S. (2022). Alcohol and Cannabinoids–From the Editors. Alcohol Research: Current Reviews, 42(1).

Mulligan, M. K., & Hamre, K. M. (2023). Influence of prenatal cannabinoid exposure on early development and beyond. Advances in Drug and Alcohol Research, 3, 10981.

Nashed, M. G., Hardy, D. B., & Laviolette, S. R. (2021). Prenatal cannabinoid exposure: emerging evidence of physiological and neuropsychiatric abnormalities. Frontiers in psychiatry, 11, 624275.

Psychoyos, D., Hungund, B., Cooper, T., & Finnell, R. H. (2008). A cannabinoid analogue of Δ9_-_tetrahydrocannabinol disrupts neural development in chick. Birth Defects Research Part B: Developmental and Reproductive Toxicology, 83(5), 477–488.

Radhakrishnan, R., Wilkinson, S. T., & D’Souza, D. C. (2014). Gone to pot–a review of the association between cannabis and psychosis. Frontiers in psychiatry, 5, 54.

Rice, J. P., Suggs, L. E., Lusk, A. V., Parker, M. O., Candelaria-Cook, F. T., Akers, K. G., Savage, D. D., & Hamilton, D. A. (2012). Effects of exposure to moderate levels of ethanol during prenatal brain development on dendritic length, branching, and spine density in the nucleus accumbens and dorsal striatum of adult rats. Alcohol, 46(6), 577–584.

Richardson, G. A., Ryan, C., Willford, J., Day, N. L., & Goldschmidt, L. (2002). Prenatal alcohol and marijuana exposure: effects on neuropsychological outcomes at 10 years. Neurotoxicology and teratology, 24(3), 309–320.

Rouzer, S. K., Cole, J. M., Johnson, J. M., Varlinskaya, E. I., & Diaz, M. R. (2017). Moderate Maternal Alcohol Exposure on Gestational Day 12 Impacts Anxiety-Like Behavior in Offspring. Front Behav Neurosci, 11, 183. doi:10.3389/fnbeh.2017.00183

Rouzer, S. K., Gutierrez, J., Larin, K. V., & Miranda, R. C. (2023). Alcohol & cannabinoid co-use: Implications for impaired fetal brain development following gestational exposure. Experimental Neurology, 114318.

SAMHSA. (2019). Mental Health Services Administration. Key substance use and mental health indicators in the United States: Results from the 2018 National Survey on Drug Use and Health (HHS Publication No. PEP19-5068, NSDUH Series H-54). Rockville, MD: Center for Behavioral Health Statistics and Quality. Substance Abuse and Mental Health Services Administration.

Serra, V., Aroni, S., Bortolato, M., Frau, R., & Melis, M. (2023). Endocannabinoid-dependent decrease of GABAergic transmission on dopaminergic neurons is associated with susceptibility to cocaine stimulant effects in pre-adolescent male MAOA hypomorphic mice exposed to early life stress. Neuropharmacology, 233, 109548. doi:10.1016/j.neuropharm.2023.109548

Smith, A. M., Fried, P. A., Hogan, M. J., & Cameron, I. (2006). Effects of prenatal marijuana on visuospatial working memory: an fMRI study in young adults. Neurotoxicol Teratol, 28(2), 286–295. doi:10.1016/j.ntt.2005.12.008

Tait, R. J., Caldicott, D., Mountain, D., Hill, S. L., & Lenton, S. (2016). A systematic review of adverse events arising from the use of synthetic cannabinoids and their associated treatment. Clinical toxicology, 54(1), 1–13.

van Amsterdam, J., Brunt, T., & van den Brink, W. (2015). The adverse health effects of synthetic cannabinoids with emphasis on psychosis-like effects. J Psychopharmacol, 29(3), 254–263. doi:10.1177/0269881114565142

Weile, L. K. K., Wu, C., Hegaard, H. K., Kesmodel, U. S., Henriksen, T. B., & Nohr, E. A. (2020). Alcohol Intake in Early Pregnancy and Risk of Attention-Deficit/Hyperactivity Disorder in Children Up to 19 Years of Age: A Cohort Study. Alcohol Clin Exp Res, 44(1), 168–177. doi:10.1111/acer.14243

Weiss, S. J., Jonn_-_Seed, M. S., & Harris_-_Muchell, C. (2007). The contribution of fetal drug exposure to temperament: potential teratogenic effects on neuropsychiatric risk. Journal of Child Psychology and Psychiatry, 48(8), 773–784.

Wieczorek, L., Fish, E. W., O’Leary-Moore, S. K., Parnell, S. E., & Sulik, K. K. (2015). Hypothalamic-pituitary-adrenal axis and behavioral dysfunction following early binge-like prenatal alcohol exposure in mice. Alcohol, 49(3), 207–217. doi:10.1016/j.alcohol.2015.01.005

Yurasek, A. M., Aston, E. R., & Metrik, J. (2017). Co-use of alcohol and cannabis: A review. Current Addiction Reports, 4, 184–193.

Zamberletti, E., & Rubino, T. (2022). Dos(e)Age: Role of Dose and Age in the Long-Term Effect of Cannabinoids on Cognition. Molecules, 27(4). doi:10.3390/molecules27041411

