## Supplementary material for "The impact of prenatal alcohol and synthetic cannabinoid exposure on behavioral adaptations in adolescent offspring and alcohol self-administration in adulthood": Figure Captions

**Figure 1.** Illustration of prenatal treatment protocol and experimental timeline.

**Figure 2.** Gestational drug exposure produces changes in baseline corticosterone (CORT) levels in male and female rats. Male **(A)** and female **(B)** rats exposed to prenatal alcohol exposure (PAE) showed a reduction in basal CORT levels. Mean ± SEM. * main effect of session, ^ main effect of PAE, $ main effect of PCE, & significant session x PAE interaction, # significant session x PCE interaction, + significant PAE x PCE interaction, † significant time x PAE x PCE interaction, p < 0.05.

/

**Figure 3.** The effects of gestational drug exposure on locomotor behavior and anxiety-like behavior in male and female rat offspring. **(A)** Male rats treated with prenatal cannabinoid exposure (PCE) - only showed reduced locomotor activity compared to rats treated with combination PAC + PCE. **(B)** There were no changes in locomotor behavior in female rats exposed to PAE or PCE. (**C**) Male rats exposed to PAE showed a general increase in % time spent in the light side of the light compartment during the light dark (LD) test. **(D)** During LD test in females, there were no significant main effects of PAE or PCE. Mean ± SEM. * main effect of session, ^ main effect of PAE, $ main effect of PCE, & significant session x PAE interaction, # significant session x PCE interaction, + significant PAE x PCE interaction, † significant time x PAE x PCE interaction, p < 0.05.

**Figure 4.** Gestational drug exposure produces differences in alcohol intake during adolescence. Alcohol intake (g/kg) across two-bottle choice sessions (3d avg) in males **(A)** and females **(C)**. Total alcohol intake (g/kg) during two-bottle choice in males **(B)** and females **(D).** Both males and female treated with PCE showed increases in total alcohol intake during two-bottle choice paradigm. Mean ± SEM. * main effect of session, ^ main effect of PAE, $ main effect of PCE, & significant session x PAE interaction, # significant session x PCE interaction, + significant PAE x PCE interaction, † significant time x PAE x PCE interaction, p < 0.05.

**Figure 5.** Gestational drug exposure produces moderate changes in alcohol intake in adult females but not males. **(A)** Male rats treated with PCE showed a general reduction in lever responses across alcohol self-administration sessions, but this was not found in total alcohol intake (g/kg) **(B).** Female rats treated with PCE showed a general increase in active lever responses across alcohol self-administration sessions **(C)** and this was also shown in total alcohol intake during adulthood **(D)**. Mean ± SEM. * main effect of session, ^ main effect of PAE, $ main effect of PCE, & significant session x PAE interaction, # significant session x PCE interaction, + significant PAE x PCE interaction, † significant time x PAE x PCE interaction, p < 0.05.

**Figure 6.** Change in reinforcer produces altered sensitivity in males but not females. Active lever responses for last two 20% EtOH sessions during alcohol acquisition session in male rats **(A)** and female **(D)** rats. Active lever responses for 1% sucrose solution in male **(B)** and female **(E)** rats. Active lever responses for 20% EtOH, 1% sucrose solution in male **(C)** and female **(F)** rats. Mean ± SEM. * main effect of session, ^ main effect of PAE, $ main effect of PCE, & significant session x PAE interaction, # significant session x PCE interaction, + significant PAE x PCE interaction, † significant time x PAE x PCE interaction, p < 0.05.

**Figure 7**. Reinitiation of drinking after abstinence period. Active lever responses during an alcohol self-administration session after a two week abstinence period in male **(A)** and female **(B)** rats. Mean ± SEM. * main effect of session, ^ main effect of PAE, $ main effect of PCE, & significant session x PAE interaction, # significant session x PCE interaction, + significant PAE x PCE interaction, † significant time x PAE x PCE interaction, p < 0.05.
